# The Social Genome of Friends and Schoolmates in the National Longitudinal Study of Adolescent to Adult Health

**DOI:** 10.1101/107045

**Authors:** Benjamin W. Domingue, Daniel W. Belsky, Jason M. Fletcher, Dalton Conley, Jason D. Boardman, Kathleen Mullan Harris

## Abstract

Humans tend to form social relationships with others who resemble them. Whether this sorting of like with like arises from historical patterns of migration, meso-level social structures in modern society, or individual-level selection of similar peers remains unsettled. Recent research has evaluated the possibility that unobserved genotypes may play an important role in the creation of homophilous relationships. We extend this work by using data from 9,500 adolescents from the National Longitudinal Study of Adolescent to Adult Health (Add Health) to examine genetic similarities among pairs of friends. While there is some evidence that friends have correlated genotypes, both at the whole-genome level as well as at trait-associated loci (via polygenic scores), further analysis suggests that meso-level forces, such as school assignment, are a principal source of genetic similarity between friends. We also observe apparent social-genetic effects in which polygenic scores of an individual’s friends and schoolmates predict the individual’s own educational attainment. In contrast, an individual’s height is unassociated with the height genetics of peers.

**Significance:** Our study reported significant findings of a “social genome” that can be quantified and studied to understand human health and behavior. In a national sample of more than 9,000 American adolescents, we found evidence of social forces that act to make friends and schoolmates more genetically similar to one another as compared to random pairs of unrelated individuals. This subtle genetic similarity was observed across the entire genome and at sets of genomic locations linked with specific traits—educational attainment and body-mass index—a phenomenon we term “social-genetic correlation.” We also find evidence of a “social-genetic effect” such that the genetics of a person’s friends and schoolmates influenced their own education, even after accounting for the person’s own genetics.

## Introduction

The degree to which genetics are implicated in the formation and consequences of social relationships is of growing interest to the new field of sociogenomics (1,2). Analysis of spousal genotypes suggests that spouses are more genetically similar to one another as compared to random pairs of individuals in the population (3–9). The degree of this genetic “homogamy” is modest. In previous analyses, we estimated that genetic homogamy was about one-third the magnitude of educational homogamy (3), even when specifically examining education-associated genotypes (8). However, even modest genetic homogamy can have implications for statistical and medical genetic models of inheritance and social models of spousal effects (10–12).

Marriage is not the only social grouping to evidence genetic selection. Adult friends are, on average, more genetically similar than random pairs from the population (13). Genetic similarity among friendship networks is important for at least two reasons. First, social networks can influence mating markets, so genetic similarity among friends may be one source of genetic similarity among spouses. Second, there may exist social-genetic effects—the effects of alter’s genotype on ego’s phenotype (1,14,15)—which would further suggest that social sorting on genotype may have consequences for the distribution of phenotypes in a population beyond its effect on subsequent generations through assortative mating.

Adolescence is a critical developmental period in which patterns of health behaviors and overall mental health established during this phase continue through the lifecourse (16) and may affect socioeconomic attainment (17,18). Moreover, it is also a time of heightened salience for peer networks and influence (19–22). For these reasons, in the present study we characterize genetic homophily within adolescent social networks in the United States. Specifically, we analyze data from the National Longitudinal Study of Adolescent to Adult Health, or Add Health (23). Add Health surveyed 90,118 US adolescents aged 12-18 in 1994-1995 using a school-based sampling frame. As part of the survey, students were asked to list the names of their friends. Responses were collated within schools to identify social ties between individuals and their friends (24). Of the adolescents surveyed, 20,745 were enrolled in a longitudinal study that included in-home interviews with the adolescents and their parents and that followed the adolescents prospectively across four waves of interviews spanning 14 years. At the most recent interview in 2008, ~12,000 Add Health participants provided DNA for genotyping and genome-wide single-nucleotide polymorphism (SNP) data were assayed. We linked these genetic data with social network information from the original school-based surveys along with information about personal characteristics and social environments accumulated across Add Health followup waves. Analyses focus on a group of genetically homogeneous respondents identified as being of ancestral European origin (N~5,500). Complete details on the data are in Section 1 of the Supplemental Information (SI).

Our analysis proceeds in three steps. In Step 1, we test if friends are more genetically similar to one another than to randomly selected peers in the same context. In Step 2, we ask about the role of school assignment in observed genetic similarity among friends. In Step 3, we evaluate a potential implication of genetic similarity among friends: social-genetic effects, or the association between the genotypes of one’s social peers and one’s own phenotype (net of own genotype).

## Results

### 1. Are friends more genetically similar to one another than they are to randomly selected peers?

We tested if friends were more genetically similar to one another as compared to random pairs of individuals. Using the KING algorithm (25), we computed genetic kinships between all pairs of Add Health participants. We then compared kinships among friends to kinships among random pairs of individuals to estimate the degree of genetic similarity among friends (3). To address potential confounding of analysis by ancestry, we focused on unrelated respondents of European ancestry. We also conducted analysis of genetic relatedness of the full Add Health sample using the REAP algorithm (26), which is designed to estimate genetic similarity in the presence of population stratification. REAP results were similar to the KING results and are reported in the SI.

Estimates of genetic similarity among friends were positive (Table 1). Among non-Hispanic whites in Add Health, KING estimates of genetic homophily were about 2/3 the magnitude of our previous KING estimates of genetic similarity among spouses in the US Health and Retirement Study (friend similarity=0.031, CI=0.022-0.036, as compared to spousal similarity=0.045 from (3)). REAP estimates of genetic similarity were somewhat smaller.

**Table 1.** Similarity estimates based on overall estimates of genetic similarity (KING) for friends and schoolmate pairs.

| Social Linkage | Friends | Schoolmates |
| --- | --- | --- |
| Relatedness Measure | KING | KING |
| N individuals | 2888 | 5611 |
| N socially-linked dyads | 4574 | 262027 |
| N possible dyads | 4168562 | 15738221 |
| Genetic Similarity | 0.031 | 0.019 |
| 95% CI | 0.022-0.036 | 0.018-0.020 |

Our second analysis tested if friends were more similar to one another on specific phenotype-related genetic dimensions. We considered the genetics of three phenotypes: height, body-mass index (BMI, a measure of adiposity), and educational attainment. We used polygenic scores to summarize phenotype-related genetics. Polygenic scores are genome-wide summaries of genetic influence. They are computed by weighting alleles at loci across the genome according to their association with a phenotype of interest and then summing weighted allele counts across loci. We computed polygenic scores based on weights from published genomewide association studies (GWAS) (27–29) using established methods (30). Because the original GWAS were performed on non-Hispanic whites, we restricted polygenic score analysis to this population as population stratification may dilute genetic associations (31). To correct for any residual population stratification, we adjusted polygenic score analyses for the first 10 principal components (32) computed based on the genetically homogeneous set of European-ancestry respondents. Further details are reported in the SI (Section 1).

We first confirmed polygenic scores were associated with their respective phenotypes (all polygenic score-phenotype correlations exceeded 0.25, Table 2). Educational attainment was strongly correlated between friends (r=0.42) and less so for BMI and height (r=0.12 for BMI, r=0.09 for height). Next, for each polygenic score, we computed associations of the respondent’s polygenic score with the average polygenic score among their friends. Polygenic scores for body mass index (r=0.08, p<0.001) and educational attainment (r=0.09, p<0.001) were positively correlated among friends. Polygenic scores for height were not correlated among friends (r=-0.01, p=0.63). As compared to correlations of polygenic scores amongst spouses (8), spousal correlations for polygenic scores for height were much larger than that observed for friends, spousal correlations of BMI polygenic scores were smaller, and friend and spousal correlations on polygenic scores for educational attainment were comparable.

**Table 2.** Correlations between various genotypes and phenotypes for individuals (as well as school ICCs) as specified in each row for the traits specified in each column. Correlations also shown for female and male respondents in which the focal individuals and the individuals used to construct measures of the social genome were restricted to the same sex. Note that not all respondents had a genotyped friend; correlations based on friend PGS are computed using a reduced sample.

|  |  | Education | BMI | Height |
| --- | --- | --- | --- | --- |
| All (n=5626) | r(phenotype,PGS) | 0.263 | 0.266 | 0.317 |
|  | Phenotype School ICC | 0.166 | 0.025 | 0.022 |
|  | r(Phenotype, Friend Phenotype) | 0.415 | 0.118 | 0.090 |
|  | r(Phenotype, School Phenotype) | 0.347 | 0.107 | 0.096 |
|  | **PGS School ICC** | **0.037** | **0.005** | **0.016** |
|  | **r(PGS, Friend PGS)** | **0.088** | **0.084** | **-0.009** |
|  | **r(PGS, School PGS)** | **0.140** | **0.042** | **0.078** |
| Females (n=2979) | r(phenotype,PGS) | 0.262 | 0.260 | 0.301 |
|  | Phenotype School ICC | 0.155 | 0.034 | 0.032 |
|  | r(Phenotype, Friend Phenotype) | 0.413 | 0.152 | 0.055 |
|  | r(Phenotype, School Phenotype) | 0.328 | 0.107 | 0.122 |
|  | PGS School ICC | 0.030 | 0.003 | 0.022 |
|  | r(PGS, Friend PGS) | 0.123 | 0.077 | 0.026 |
|  | r(PGS, School PGS) | 0.110 | 0.026 | 0.077 |
| Males (n=2647) | r(phenotype,PGS) | 0.264 | 0.276 | 0.336 |
|  | Phenotype School ICC | 0.147 | 0.009 | 0.028 |
|  | r(Phenotype, Friend Phenotype) | 0.371 | 0.119 | 0.150 |
|  | r(Phenotype, School Phenotype) | 0.318 | 0.031 | 0.086 |
|  | PGS School ICC | 0.040 | 0.005 | 0.012 |
|  | r(PGS, Friend PGS) | 0.097 | 0.108 | -0.003 |
|  | r(PGS, School PGS) | 0.133 | 0.029 | 0.033 |

Research suggests that there may be important differences among men and women with respect to the role of genes in complex social behaviors (33). To test for such differences, we repeated our analysis within same-sex social networks; that is, we tested correlations between women’s polygenic scores and scores of their female friends and between men’s polygenic scores and scores of their male friends. Results were similar to the mixed-sex analysis (Table 2). For education polygenic scores, women’s female social networks demonstrated modestly stronger social-genetic correlations as compared to men’s male social networks. The opposite was observed for BMI polygenic scores.

### 2. Why are friends more genetically similar to one another than they are to randomly selected peers?

We considered two hypotheses. One hypothesis is that friends are more genetically similar to one another because they form their friendships partly on the basis of shared characteristics (e.g. being short or tall, heavy or slim, from well-educated or poorly-educated families, etc.). This process is called “social homophily” (34–37). When characteristics that influence formation of social ties are heritable, which many are (38), social homophily can generate genetic similarity between friends. A second hypothesis is that friends are more genetically similar because people tend to form friendships within environments that are socially stratified (e.g., living in the same community, attending the same school). We refer to this process, which has been observed as a cause of demographic similarity among spouses (39), as social structuring (40). When genetics influence the social environments people live in—e.g. through influence on socioeconomic attainment (41)—social structuring can generate genetic similarity between friends even without explicit selection on phenotypic similarity. Social homophily and social structuring are not mutually exclusive and may indeed be complementary processes.

Our analysis of polygenic score correlations among friends suggested some evidence of social homophily; individuals were modestly similar to friends in terms of their educational trajectories (measured as their educational attainment at Wave IV, about 15 years after social network data were originally collected) and their educational attainment polygenic scores were correlated (Table 2).

We next tested for evidence of social structuring of genetic similarity. We estimated genetic similarity at the level of a structural social environment—the schools within which social networks of friends were defined. For this analysis, we compared genetic similarity among schoolmates to genetic similarity among random pairs of individuals in the Add Health sample. Schoolmates were genetically more similar to one another as compared to random pairs of individuals (for European schoolmates, KING similarity=0.019, CI=0.018-0.020; Table 1).

Given this evidence of social structuring, we repeated analysis of genetic similarity between friends, this time comparing friends to random pairs of individuals drawn from the same school. Genetic similarity between friends was attenuated by about half when the analysis compared friends to random pairs of schoolmates (KING-estimated within-school friend similarity=0.018, CI=0.009-0.026; Figure 1A). In parallel, correlations among educational attainment polygenic scores between friends were attenuated when we adjusted analysis for the school mean polygenic score (Figure 1B; underlying coefficients are reported in SI Section 3); less attenuation was observed for estimates related to height and BMI polygenic scores.

**Figure 1.**
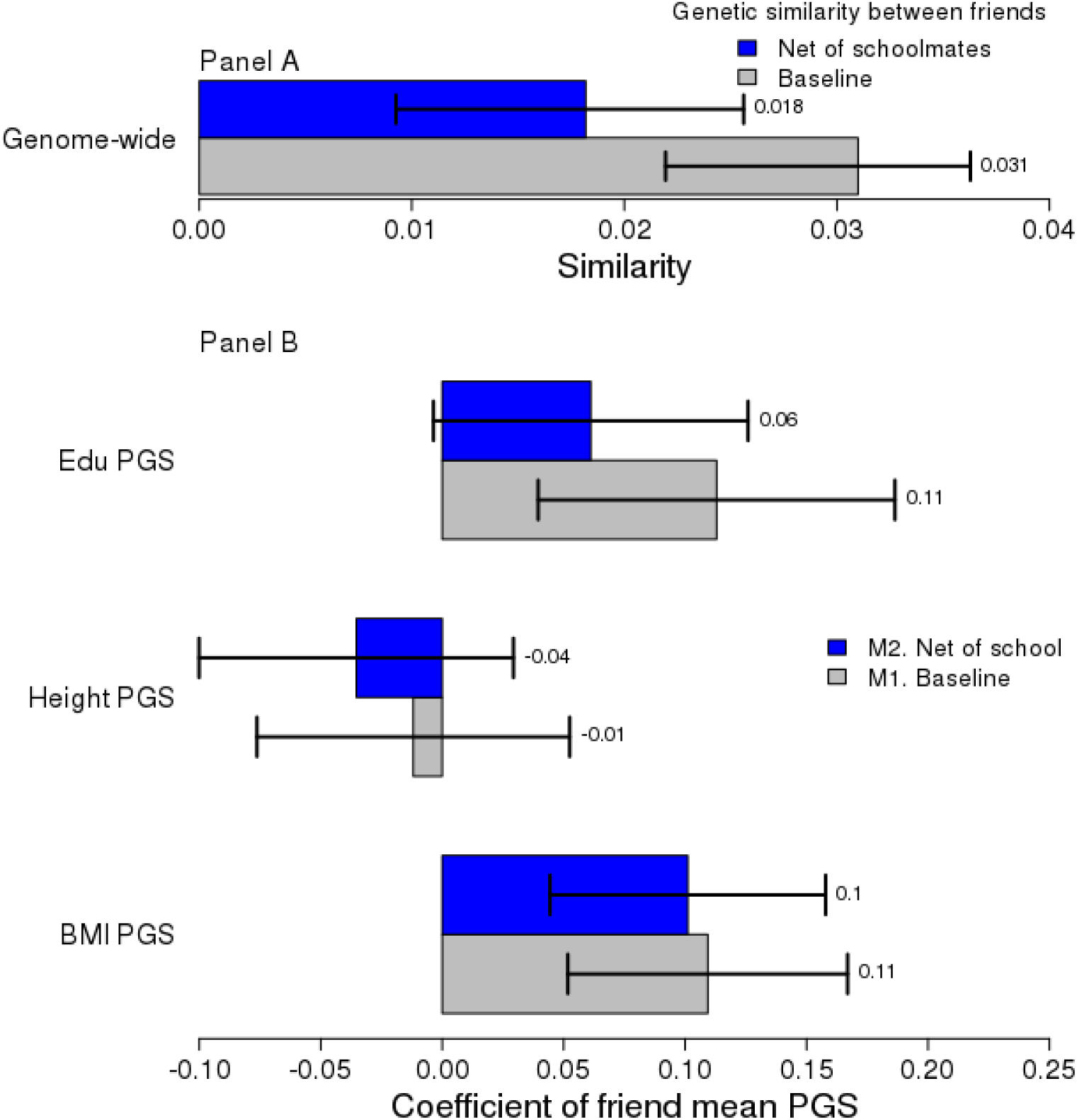
Social-genetic correlations: Panel A. Genetic similarity between friends before and after accounting for genetic similarity of schoolmates. Panel B. Associations between participants’ polygenic scores and the polygenic scores of their friends before (M1) and after (M2) adjusting for the mean polygenic score for participants’ school. CIs are robust to school clustering.

### 3. Is the social genome associated with an individual’s phenotype?

One reason genetic similarity among friends and schoolmates may matter for health and social science research is the potential phenomenon of social-genetic effects. Social-genetic effects, also called “indirect” genetic effects, refer to the influence of one organism’s genotype on a different organism’s phenotype (14,15). Social-genetic effects, which may take several forms (1), are accounted for in evolutionary theory (42,43) and have been observed among animals (15,44–47) and received some study in human siblings (14,48,49).

We tested social-genetic effects among unrelated individuals with direct social ties (friends) and structural social ties (schoolmates). We examined two types of social-genetic effects: (a) social-genetic main effects—associations between friend or school genetics and a focal individual’s phenotype net of that focal individual’s own genetics; and (b) social epistasis— moderation of the association between a focal individual’s own genetics and phenotype by the genetics of their social environment. This analysis comes with the caveat that social ties between friends are not randomly assigned and thus estimates of social-genetic effects cannot be strictly interpreted as causal.

We tested social-genetic main effects among friends by analyzing associations between average friend polygenic scores and a focal Add Health participant’s educational attainment, body-mass index, and height measured at the Wave IV Add Health follow-up, 13 years after social network information was collected. Friend polygenic scores for educational attainment were associated with Add Health participants’ educational attainment at the Wave IV assessment (b=0.18, p<0.001). This apparent social-genetic effect was not explained by social-genetic correlations among friends; when the focal individual’s own education polygenic score was included as a covariate, the social-genetic effect was only slightly reduced and remained statistically significant (b=0.15, p<0.001). No friend-level social-genetic main effects were observed for body-mass index or height (SI section 4). Given that weight may be more sensitive to context than height (e.g., a friend’s appetite or interest in exercise may potentially influence a focal individual’s weight but presumably not their height), we also considered an alternative adiposity measure (SI Section 4). Results were similar.

We next tested social-genetic main effects at the school level after accounting for the individual’s own polygenic score. Findings were similar to findings for friends. Attending a school with higher average education polygenic score predicted completing more years of schooling by the time of Wave IV follow-up 13 years later (after adjustment for the person’s own polygenic score, b=0.22, p<0.001). We also observed a weak school-level social-genetic effect for body-mass index (SI section 4). Analyses of social-genetic effects are summarized in Figure 2 (underlying coefficients are reported in SI Section 4).

**Figure 2.**
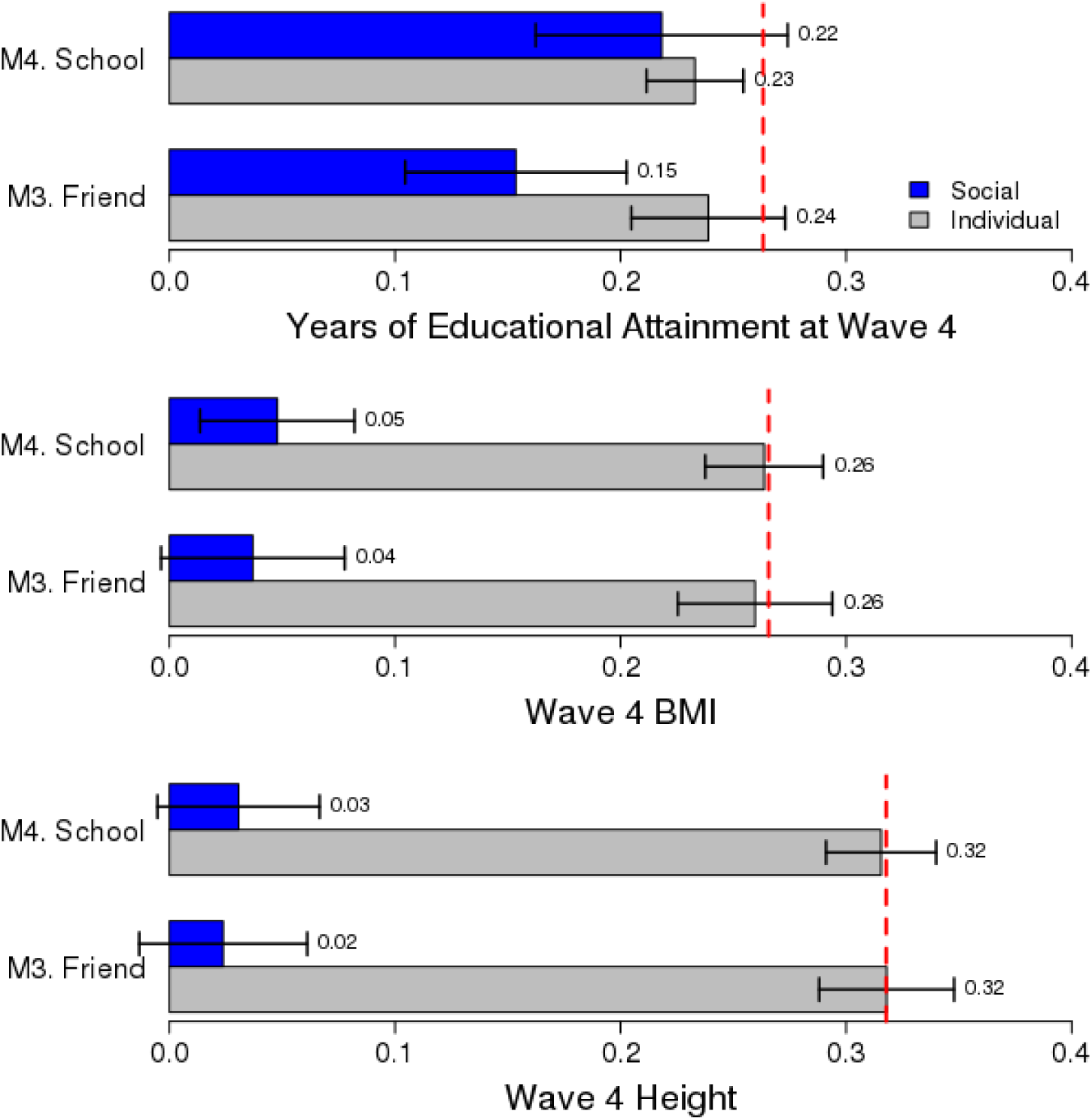
Social-genetic Effects: Effect of friend and school mean PGS net of one’s own PGS for educational attainment, BMI, and height (all measured at Wave 4) on associated outcome. Outcomes are standardized as are social genotypes. The red line is the baseline effect of own PGS on the outcome in a null model with no other predictors. Estimates are based on a sample of unrelated respondents. CIs are robust to school clustering.

To test if school-level social-genetic main effects might account for social-genetic main effects among friends, we re-estimated friend-level social-genetic main effects using a within-school design. This analysis focuses comparisons between individuals within a school through inclusion of a school fixed effect. For educational attainment, friend-level social-genetic main effects remained statistically significant even when comparing an individual to their schoolmates (within-school social-genetic main effect b=0.1, p<0.001). We conducted sensitivity analyses to evaluate whether findings were affected by genetic relatedness among groups of schoolmates and friends (7), by particular schools that included large numbers of respondents, by the strength of social ties between friends (mutual nominations; first and second degree friendship networks), or by the ethnic composition of the schools. Results were similar to those reported (SI section 4).

Finally, we tested for social epistasis by analyzing moderation of the association between Add Health participant’s own polygenic scores and their phenotypes by friend- and schoolmate-average polygenic scores. We found some evidence of social epistatic effects predicting educational attainment. The association between a respondent’s polygenic score and their own educational attainment was of somewhat larger magnitude when their friends and schoolmates had higher average educational attainment polygenic scores (Section 4 of SI).

## Discussion

We analyzed genome-wide and social network data from a large nationally-representative sample of American adolescents in the National Longitudinal Study of Adolescent to Adult Health. There were three main findings. First, we found evidence for positive genetic similarity amongst friends; friendship pairs tended to be more genetically similar to one another compared to random pairs of individuals. Second, friends tend to be genetically similar because of two potentially complementary processes, social homophily and social structuring. Social homophily-related genetic similarity may arise because individuals form social bonds on the basis of characteristics they share, many of which have genetic origins. In Add Health, friends tended to share educational trajectories and also to have similar education-associated genotypes, suggesting social-homophily-related genetic similarity. Social structuring-related genetic similarity may arise because individuals tend to form social bonds with people whom they share the same social environments and spend their time together and these social environments may be partly influenced by their genetics. Genetic similarity at the structural level, i.e. among schoolmates, accounted for about half of the genetic similarity among the Add Health friends. Third, genetic similarity of friends and schoolmates may bias interpretations of genetic main effects. We found evidence that genetics tended to be somewhat more similar among socially connected individuals and that the genetics of individuals in a person’s social environment influence that person’s phenotype. Together, these findings imply that naïve estimates of genetic associations that ignore the social genome may be modestly biased by unmeasured social-genetic effects. In our analysis of education-associated genetics, the direction of this bias was positive. For other phenotypes and in other social settings, bias may be negative.

### Implications for Genetics

Our findings regarding genetic similarity among friends echo and extend earlier work suggesting that adult friends exhibit overall genetic similarity (13); adolescents exhibit similarity on specific candidate genes (50,51); and genetic similarities between adolescent friends may be partially responsible for observed similarities of friend GPAs (52). Although power to detect genome-wide genetic similarity in our analysis may be low (7), we observed consistent evidence in analysis of polygenic scores. Potential confounding by population stratification is also an issue (7,53). However, our results were consistent when we analyzed genetic similarity using the REAP software, which is designed to account for population stratification (26) and when we analyzed increasingly homogeneous subsamples.

A novel observation from our analysis is that genetic similarity among persons with direct social ties (friends) partly reflects genetic similarity among persons with structural social ties (schoolmates). This school-level similarity was also observed at the level of trait-associated genetics. About 4% of variation in the educational attainment polygenic score was between schools. Future research into social-genetics in humans should be designed to account for this structural dimension, or other potential ways in which individuals may be clustered (54). Of course, we only measured certain aspects of social structuring. Studies that probe whether the magnitude of social structuring—perhaps as proxied by differences in institutional features (e.g., public versus private schools; school-level segregation (51)) or even geographic segregation (55)—drives the degree of genetic assortment would also be informative.

A second novel observation from our analysis is that the genetics of a person’s social network may affect that person’s risk of obesity and their educational attainment. As noted above, such social-genetic effects can bias analysis of genetic associations if the social genome is not accounted for (15). It is noteworthy that our social-genetic effect results were specific to phenotypes plausibly influenced by the social environment, education and obesity (see also (56,57)). For height, which is not likely to be influenced by social network processes (56), at least in contemporary US society, we observed no evidence of social-genetic effects. Future research may begin to bridge the social genetic effects literature with that of peer- and school-effects by examining whether observable characteristics of one’s social environment (e.g., the psychological makeup of one’s friends, their substance abuse behavior, personality traits, etc.) act as mechanisms through which social genetic effects occur.

### Implications for Social Science

Putative social-genetic effects also have implications for social science research. Because our analysis is able to control for the same genetics in the adolescents that are measured in their friends and schoolmates, the social-genetic effects we detect are evidence of environmental transmission of peer- and school-level influences on adolescents’ outcomes. A critical next step is to determine whether social-genetic effects detected in analysis of friends and schoolmates reflect causal effects arising from friends and schoolmates, or if the genetics of friends and schoolmates function as proxies for other features of adolescents’ environments. For example, in the case of education, friend- and school-level genetics may be associated with other features of communities that promote student attainment (58). In order to be more definitive, future research may require exogenous mechanisms underlying social contact (59,60). Specifically, others have argued that larger social environments such as schools are excellent environments for gene-environment research because school selection is largely independent of genotype (61) but our results suggest that this may not be accurate.

To the extent social-genetic effects are causal, they provide new opportunities to investigate social network processes. Many previous attempts to identify causal social-network effects (62) have been met with skepticism (56). In observational studies of friends, one of the fundamental problem is that the friendship bond is endogenous to the characteristics of the individuals in the friendship, making it challenging to disentangle the effects of friends from effects of characteristics contributing to friendship formation (63). A second, equally daunting challenge is the “reflection problem” (64) wherein it is difficult to distinguish the direction of effects in social interaction. Because a person’s genetics cannot be caused by social processes occurring within that person’s lifetime, they can help distinguish homophily (like assorting with like) from contagion in social network research. To the extent that genotype and environments are independent and randomly assigned, such social-genetic analysis can also solve the reflection problem—a strong assumption, however (65).

This study contributes to what is currently known about the role of genotypes with respect to the social ecology that exists among humans. We have provided specific evidence about genetic similarity within social networks and the potential for social-genetic effects. The joint existence of social-network genetic similarity and social-genetic effects could produce important feedback loops. If the genetics of one’s social environment matter and relevant genetics are stratified across environments, then being in certain social environments might confound straightforward analysis of genetic effects (15). More work is needed to document these features of social genetics in humans before such modulations can be unambiguously documented, but their possible existence challenges simple notions of genetic “effects.”

## Methods & Materials

### Data

The National Longitudinal Study of Adolescent to Adult Health (Add Health) is a nationally representative cohort drawn from a probability sample of 80 US high schools and 52 US middle schools (in 80 US communities), representative of US schools in 1994–95 with respect to region, urban setting, school size, school type, and race or ethnic background (22,23). About 15,000 respondents (or 96%) consented to genotyping during the Wave 4 interview in 2008–09 for purposes of approved Add Health Wave 4 research. Of those who consented to genotyping, ~12,000 (or 80%) agreed to have their DNA archived for future testing (see SI for a comparison of genotyped and non-genotyped respondents). DNA extraction and genotyping on this archived sample yielded a sample of 9,975 Add Health members with GWAS data consisting of 631,990 SNPs. The SI contains additional details on the genotyping process.

In the in-school survey (which was administered to every student in the participating schools, not only the Add Health study members who are prospectively followed into adulthood), as well as in the in-home surveys at waves 1 and 2, students were asked to nominate up to 5 of their male friends and 5 female friends. We accept a nomination in either direction (i.e. “undirected” friendships) as evidence of a friendship between two individuals. Of those with genetic data, only 5,199 people were in a friendship pair with another genotyped respondent. We focus largely on friendship nominations within race/ethnicity. Of the 7,217 friendship pairs between genotyped respondents, ~90% were within self-reported race/ethnicity. We emphasize two additional caveats. First, related respondents (as identified by measures of genetic similarity) were not included in the analyses. Second, one school—a so called “saturated school” in the Add Health data (23)—contributed a disproportionate number of friend pairs to the sample of non-Hispanic white respondents. Results reported are robust to the removal of this school (see SI).

### Measures

To measure genetic similarity, we use a kinship measures (KING; (25)) that has been the focus of earlier research (3,5). We also consider an alternative measure (REAP (26)) that is less sensitive to population stratification. We construct principal components using all genotyped respondents. We construct polygenic scores for anthropometric traits (BMI, height) and educational attainment. To construct these scores, we utilize publicly available genome-wide association study results (27–29) derived from large consortia studies (that did not include Add Health) of populations of European descent. Polygenic scores do not typically generalize across racial groups (66), so we focus these analyses on a group of genetically homogeneous European-ancestry respondents.

We use information from Wave 4 (when respondents were 24–32 years old) on years of education completed, BMI, and height (see additional details on all measures in SI). For analysis, we first residualized all three outcomes on sex and birth year.

## Methods

### Overall Genetic Similarity

We first consider a measure of genetic similarity amongst friends used previously to study genetic similarity amongst spouses (3,5). We compute the area between the 45 degree line and the P-P plot (given two CDFs F and G, the P-P plot is the set of points (F(x),G(x)) for all x) comparing the density of genetic similarity between friends with the density of genetic similarity for all dyads.

### Targeted Genetic Similarity

We first consider a baseline model for some polygenic score G_i_ of the form

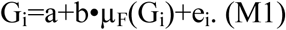

where µ_F_(G_i_) is the mean of G_i_ for all an individual’s friends. The coefficient b captures the degree to which being friends is associated with levels of overall genetic similarity. Motivated by previous studies of Add Health social networks (67,68), we next consider the role of genetic clustering into schools in the observed degree of friend genetic similarity via

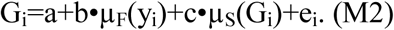

where µ_S_(G_i_) is the mean of G_i_ for all other individual’s at the school of individual i. We interpret attenuation in estimates of b going from M1 to M2 as evidence for the importance of social structure, specifically school assignment, in observed genetic similarity amongst friends. We adjust all standard errors for clustering of students into schools (69).

Turning to our analysis of social-genetic effects, we first consider models of the (potentially confounded) effect of the social genome on an individual’s phenotype (P_i_) net of the individual’s own polygenic score (PGS_i_). The social genome will be characterized via µ_F_(G_i_) and µ_S_(G_i_) as above. We first consider

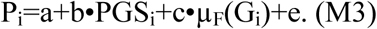

We construe this as a test of “narrow” social-genetic effects of friends. In contrast, we also consider measures of “broad” social-genetic effects:

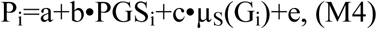

Again, all standard errors are adjusted for clustering into schools.

## Acknowledgments

This research uses data from Add Health, a program project directed by Kathleen Mullan Harris and designed by J. Richard Udry, Peter S. Bearman, and Kathleen Mullan Harris at the University of North Carolina at Chapel Hill, and funded by grant P01-HD31921 from the Eunice Kennedy Shriver National Institute of Child Health and Human Development, with cooperative funding from 23 other federal agencies and foundations. Information on how to obtain the Add Health data files is available on the Add Health website (http://www.cpc.unc.edu/addhealth). We gratefully acknowledge funding from NICHD to Boardman and Harris through grants R01HD073342 and R01HD060726. Belsky is supported by a Jacobs Foundation Early Career Research Fellowship and by NIA grants R01AG032282 and P30AG028716. Conley is supported by a Russell Sage Foundation grant on “GxE and Health Inequality across the Lifecourse” (83-15-29). This research benefitted from GWAS results made publicly available by the SSGAC and GIANT consortia. We would like to acknowledge Matthew Robinson for helpful comments and Robbee Wedow, Joyce Tabor, Heather Highland, and Christy Avery for assistance with the Add Health genetic sample.

